# Copy-number variation contributes 9% of pathogenicity in the inherited retinal degenerations

**DOI:** 10.1101/742106

**Authors:** Erin Zampaglione, Benyam Kinde, Emily M. Place, Daniel Navarro-Gomez, Matthew Maher, Farzad Jamshidi, Sherwin Nassiri, J. Alex Mazzone, Caitlin Finn, Dana Schlegel, Jason Comander, Eric A. Pierce, Kinga M. Bujakowska

**Author notes:** these authors contributed equally to the work (Erin Zampaglione, Benyam Kinde). Corresponding author: Dr. Kinga Bujakowska, Massachusetts Eye and Ear, 243 Charles Street, Boston, MA 02114.

## Abstract

**Purpose:** Current sequencing strategies can genetically solve 55-60% of inherited retinal degeneration (IRD) cases, despite recent progress in sequencing. This can partially be attributed to elusive pathogenic variants (PVs) in known IRD genes, including copy number variations (CNVs), which we believe are a major contributor to unsolved IRD cases.

**Methods:** Five hundred IRD patients were analyzed with targeted next generation sequencing (NGS). The NGS data was used to detect CNVs with ExomeDepth and gCNV and the results were compared to CNV detection with a SNP-Array. Likely causal CNV predictions were validated by quantitative (q)PCR.

**Results:** Likely disease-causing single nucleotide variants (SNVs) and small indels were found in 55.8% of subjects. PVs in *USH2A* (11.6%), *RPGR* (4%) and *EYS* (4%) were the most common. Likely causal CNVs were found in an additional 8.8% of patients. Of the three CNV detection methods, gCNV showed the highest accuracy. Approximately 30% of unsolved subjects had a single likely PV in a recessive IRD gene.

**Conclusions:** CNV detection using NGS-based algorithms is a reliable method that greatly increases the genetic diagnostic rate of IRDs. Experimentally validating CNVs helps estimate the rate at which IRDs might be solved by a CNV plus a more elusive variant.

## INTRODUCTION

Over two million people worldwide are affected by inherited retinal degenerations (IRDs), a family of blinding diseases characterized by progressive death and dysfunction of primarily rod and cone photoreceptors^1,2^. Pathogenic variants (PVs) in over 270 genes have been associated with IRDs^3^, many of which were discovered recently by virtue of advances in sequencing technologies^4–23^. However, despite substantial progress in genetic methodologies, current strategies can genetically solve only about 55-60% of IRD cases^24–38^. The remaining missing diagnoses are in part due to new, yet to be discovered IRD genes. However, PVs in each new disease gene are rare, affecting a handful of IRD patients^4–23^, suggesting that the missing genetic causality largely lies in the known IRD genes. A considerable proportion of these elusive PVs are due to structural variations (SVs) such as copy number variations (CNVs), or deep intronic variants that affect splicing^36,39–49^, which are not readily available from the standard output of targeted next generation sequencing (NGS) pipelines.

Our previous work analyzed 28 genetically unsolved families with whole exome sequencing (WES) and SNP and/or Comparative Genomic Hybridization (CGH) arrays, and showed that large deletions in known IRD genes were responsible for disease in five of the families (18% of unsolved cases)^41^. In this study we applied further bioinformatic analyses that permit detection of CNVs on the panel-based NGS Genetic Eye Disease (GEDi) diagnostic test that involves sequencing the exons of all known IRD disease genes^24,50–52^. To assess the accuracy of CNV calling based on the NGS read depth we compared two algorithms, ExomeDepth^50^ and gCNV^52^, with the SNP-array based approach^53^. A subset of the CNVs were subsequently validated by qPCR. In addition, we specifically searched for the *Alu* transposable element insertion in *MAK*, which is a common cause of IRD in people of Ashkenazi Jewish descent^54–56^. Applying these techniques improved the genetic diagnostic rate for IRD patients by 10%.

## METHODS

### Ethical guidelines

The study was approved by the institutional review board at the Massachusetts Eye and Ear (Human Studies Committee MEE in USA) and adhered to the tenets of the Declaration of Helsinki. Informed consent was obtained from all individuals on whom genetic testing and further molecular evaluations were performed.

### Clinical evaluation

Patients included in the study were recruited and clinically examined at MEE. Ophthalmic examination included best-corrected Snellen visual acuity, dynamic Goldmann visual field testing, dark adaptation testing and full-field electroretinographic (ERG) testing with assessment of 0.5 Hz ERG amplitude and 30 Hz ERG amplitudes.

### Genomic (g)DNA extraction and targeted sequencing

DNA was extracted from venous blood using the DNeasy Blood and Tissue Kit (Qiagen, Hilden, Germany). All samples underwent Genetic Eye Disease test (GEDi) sequencing as described previously^24^. The GEDi version used in this study included 266 genes known to be associated with monogenic inherited retinal degenerations^3,24^ (Supplementary Table S5). The capture libraries were sequenced on MiSeq (9 samples per run) or HiSeq (96 samples per run) NGS platforms (Illumina, San Diego, CA) as previously described^24^. The NGS data was analyzed using Genome Analysis Toolkit (GATK)^51^ version 3 and annotated using the Variant Effect Predictor (VEP) tool^57^ with additional annotations taken from the Genome Aggregation Database (GnomAD) Genomic Evolutionary Rate Profiling (GERP), SIFT, PolyPhen2 and retinal expression^58^. Rare variants were selected based on the minor allele frequency (MAF) in public databases of less than 0.3%. Variants were annotated based on the transcripts included in Supplementary Table S5.

Exon 15 of *RPGR* transcript NM_001034853 (called *RPGR* ORF15) is not fully covered by the NGS and therefore it was PCR amplified and Sanger sequenced with previously established protocols (Supplementary Methods).

### Copy Number Variation (CNV) Analysis

#### NGS read-depth Analysis

Copy number variation from NGS read-depth was inferred using ExomeDepth^50^ and gCNV from the GATK version 4^51^. Samples from all of the MiSeq runs were processed together (193 samples) and HiSeq runs consisting of 96 samples each (except for one 48 sample run) were analyzed separately. In ExomeDepth analysis the samples were separated by gender, in gCNV analysis they were kept together. In the gCNV analyses, the GEDi-captured regions were padded by 250bp on each side and they were run in COHORT mode without external control samples. CNVs present in more than 15% samples were removed as they were considered to be either capture artifacts or common CNVs that would not lead to a rare Mendelian disorder. In addition, we removed CNVs in the *OPN1LW* gene and the *OPN1MW* gene, which were likely artifacts of poor mapping quality of the NGS reads.

#### SNP-Array

Genomic DNA (gDNA) samples from probands were analyzed with whole-genome SNP microarray (HumanOmni2.5 BeadChip, Illumina) according to the manufacturer’s instructions. The hybridized SNP arrays were analyzed using an array reader (iScan array scanner, Illumina) and the SNP calls were made with the genotyping module of the data analysis software (GenomeStudio, Illumina). Copy number variation from the SNP-array results was detected with PennCNV using default parameters and sex information^53^.

### Quantitative real-time PCR

Deletions were validated using quantitative real-time PCR (qRT-PCR or qPCR) on gDNA with primers specific to sequences inside the presumed CNV and flanking the CNV (Supplementary Table S1). The amplification was normalized to the *ZNF80* reference gene. For each qPCR reaction 5 ng of gDNA, 200 nM of each primer and 10 μl of Fast SYBR Green master mix (Life Technologies, Grand Island, NY) were used. The amplification was performed in a qPCR system (Stratagene Mx3000P®, Agilent Technologies) using the standard thermo-cycling program: 95°C for 3 minutes, 40 cycles of 95°C for 20 seconds, 60°C for 1 minute followed by a melting curve. Each sample was assayed in triplicate. Relative changes in genomic sequence abundance were calculated using the 2^-ddC^_T_ method^59^, and error was calculated using standard propagation of errors. Data was visualized using custom R scripts.

## RESULTS

A cohort of 500 IRD patients was sequenced with the targeted NGS panel – Genetic Eye Disease test (GEDi)^24^ and analyzed with a standard NGS analysis pipeline detecting single nucleotide variants (SNVs) and small insertions and deletions (indels)^51^ to find likely pathogenic variants leading to retinal degeneration. SNVs and small indels likely leading to disease were found in 279 IRD subjects (55.8% of the total cohort), where PVs in *USH2A* (11.8%), *RPGR* (4%) and *EYS* (4%) were the most common causes of disease (Figure 1, Supplementary Table S2). In addition, *MAK-Alu* insertion was detected in seven patients by analyzing the NGS sequence reads^60^.

**Figure 1:**
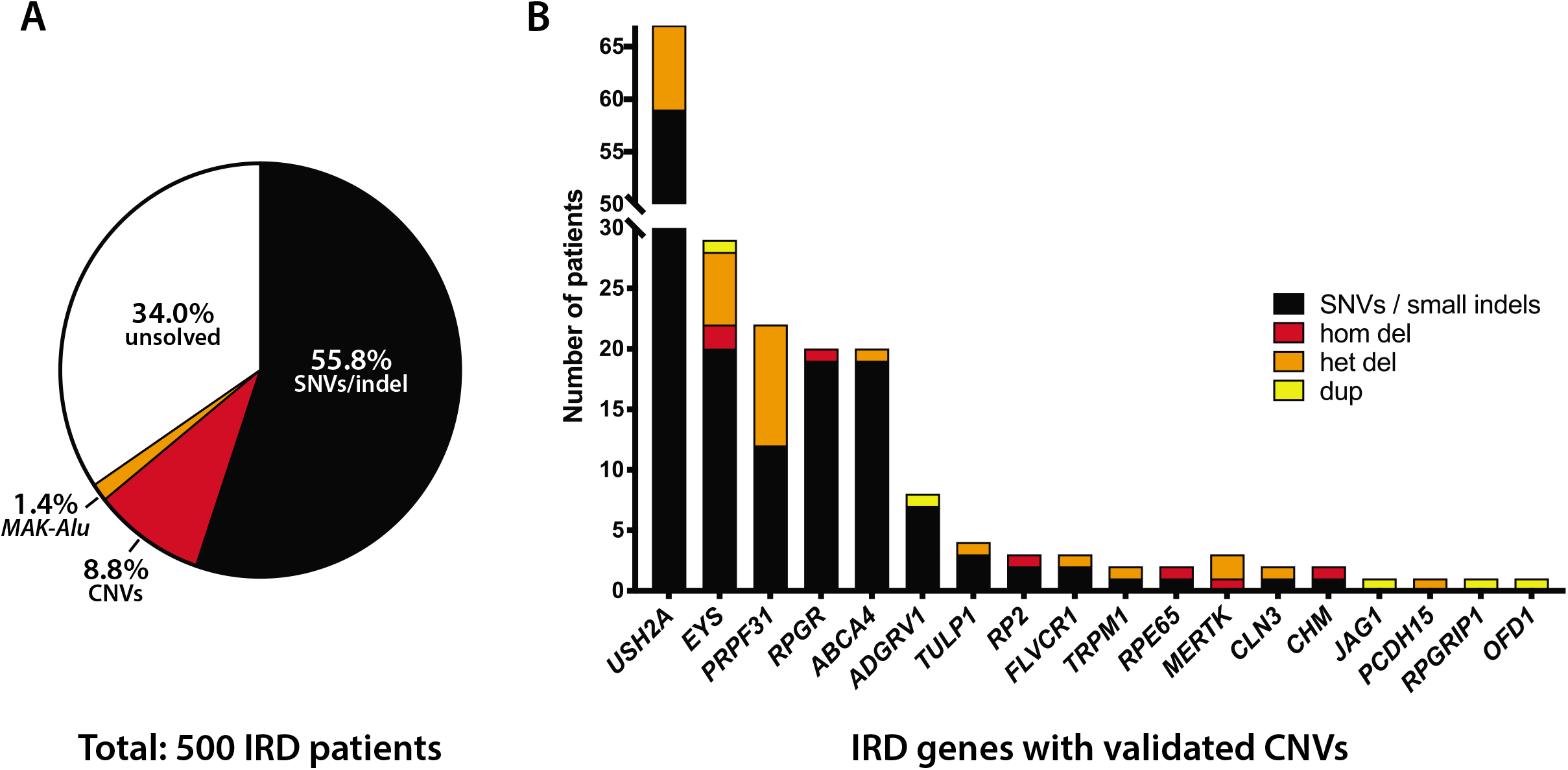
Summary of Genetic Contributions to IRD. A) Targeted NGS analysis in a cohort of 500 IRD subjects reveals SNV/small indel solutions in 55.8% cases, CNV solutions in 8.8% cases and MAK-Alu insertions in 1.4% cases. B) Breakdown of CNV solutions by gene, with the number of patients solved by SNVs in the same gene. Note that genes that commonly have SNV solutions *(USH2A, EYS)* also tend to have CNV solutions, a notable exception being *PRPF31*, in which CNVs are more common than expected based on the number of SNVs.

### CNV predictions using NGS data

Additional analysis of the NGS sequence data to detect copy number variations was performed with ExomeDepth^50^ and gCNV^52^. Our initial analysis focused on CNVs that would contribute to solving the case genetically, that is: a) homozygous CNVs in autosomal recessive (ar) genes; b) heterozygous CNVs in ar genes which are in *trans* with likely causal SNVs or small indel in the same gene; c) compound heterozygous CNVs in ar genes; d) heterozygous CNVs in a haploinsufficiency gene; e) duplications in autosomal dominant genes, which could lead to gain of function variants. The CNVs that fulfilled these criteria were subsequently validated by qPCR using genomic DNA from patients and healthy controls. This analysis revealed likely genetic solutions for an additional 44 patients (8.8%). Thirty-one of them carried heterozygous deletions, eight carried homozygous deletions and five carried heterozygous duplications. CNVs in *PRPF31* (10), *EYS* (9) and *USH2A* (8) were the most common (Figure 1B, Supplementary Table S3).

Most of the CNVs detected in this project were easily interpretable as they were either homozygous deletions, heterozygous deletions coupled with a deleterious allele in *trans* or heterozygous deletions in the known haploinsufficiency gene *PRPF31*^61^. However, there were a few examples of more unusual CNV contributions to disease etiology.

In two patients we detected two non-consecutive heterozygous deletions, which are thought to be due to deletions in *trans* (e.g. *EYS* deletions in subject 121-182 (simplex RP), or *PCDH15* deletions in subject OGI635_001299 (Usher Syndrome), Supplementary Table S3, Supplementary Figure S1). Unfortunately, due to lack of samples from family members and a large distance between the deletions (0.86 and 0. 51 Mb), the phase of these deletions could not be confirmed.

Subject OGI655_001331 presented with inherited macular degeneration, historically known as Stargardt Disease, and carried a heterozygous deletion of almost the entire *ABCA4* gene (exons 1-40). On closer examination, it was revealed that the same subject also carried a common hypomorphic missense change p.Asn1868Ile, which due to its high allele frequency (AF=0.042 in GnomAD) was overlooked during the initial analysis. The p.Asn1868Ile missense in *trans* with a deleterious missense variant or loss of function allele was shown to lead to a late onset macular degeneration^62^. Subject OGI655_001331 presented with asymptomatic macular changes at age 40 at routine eye examination, which led to the diagnosis of inherited macular degeneration/Stargardt disease at age 42, which corresponds to the late onset symptoms published by Zernant and colleagues^62^.

We identified five duplications (Figure 1B, Supplementary Table S3), two of which are highlighted below. Subject OGI2839_004424 carried a heterozygous duplication of exons 1-25 (3199 coding base-pairs) in *JAG1* gene, which causes a frameshift leading to a loss of function allele. Heterozygous loss of function variants in *JAG1* are known to lead to Alagille syndrome through haploinsufficiency of this gene^63^. The major clinical feature of Alagille syndrome are abnormalities in bile ducts leading to liver damage and accompanying features are congenital heart defects, vertebrae anomalies and ocular defects including pigmentary retinopathy and optic disc drusen^64^. Investigation of the clinical notes revealed that subject OGI2839_004424 was seen in the ophthalmic clinic for the retinopathy, but in addition was previously diagnosed with Alagille syndrome. Subject OGI2829_004414 is a male that presented with retinal degeneration and history of kidney transplantation and possible cognitive dysfunction. No variants in ciliopathy genes were identified with GEDi, however a duplication of exons 6-15 in *OFD1* was predicted by gCNV and validated by qPCR. This duplication will likely lead to an inframe duplication of 1242 internal coding bases (414 amino-acid residues), with presumed partial preservation of protein function as complete loss of function of the gene is embryonic lethal in males^65^.

### Comparison of CNVs predicted by NGS analysis versus SNP-array analysis

To assess the diagnostic performance of ExomeDepth and gCNV, we compared them with the SNP-Array-based detection of CNVs using the PennCNV algorithm^53^. One hundred and forty-four samples out of the total 500 samples from the NGS cohort were analyzed with the Illumina Omni 2.5 SNP array. In the 144 samples, there were 61 predicted solving CNVs by gCNV and ExomeDepth in 54 patients. The remaining 90 patients were chosen at random. In these 144 samples, gCNV predicted 52 potentially solving CNVs, ExomeDepth predicted 52 potentially solving CNVs, and SA predicted 31 potentially solving CNVs. All predicted solving CNVs were then tested by qPCR to determine if they were confirmed true positives (Figure 2A, B, Supplementary Figure S1A, B). Of the three methods used, gCNV showed the best performance with 44 out of 52 predictions that validated (positive predictive value, or PPV equal to 85%) compared to ExomeDepth (37/52, PPV=71%) and SNP-Array (25/31, PPV=81%) (Table 1). Intersection of any two algorithms increased the PPV with a considerable decrease of sensitivity (Table 1). The gCNV algorithm alone predicted all of the true positive CNVs detected in this study (Figure 2A). False positive rate was the highest (29%) in ExomeDepth (Figure 2B).

**Figure 2:**
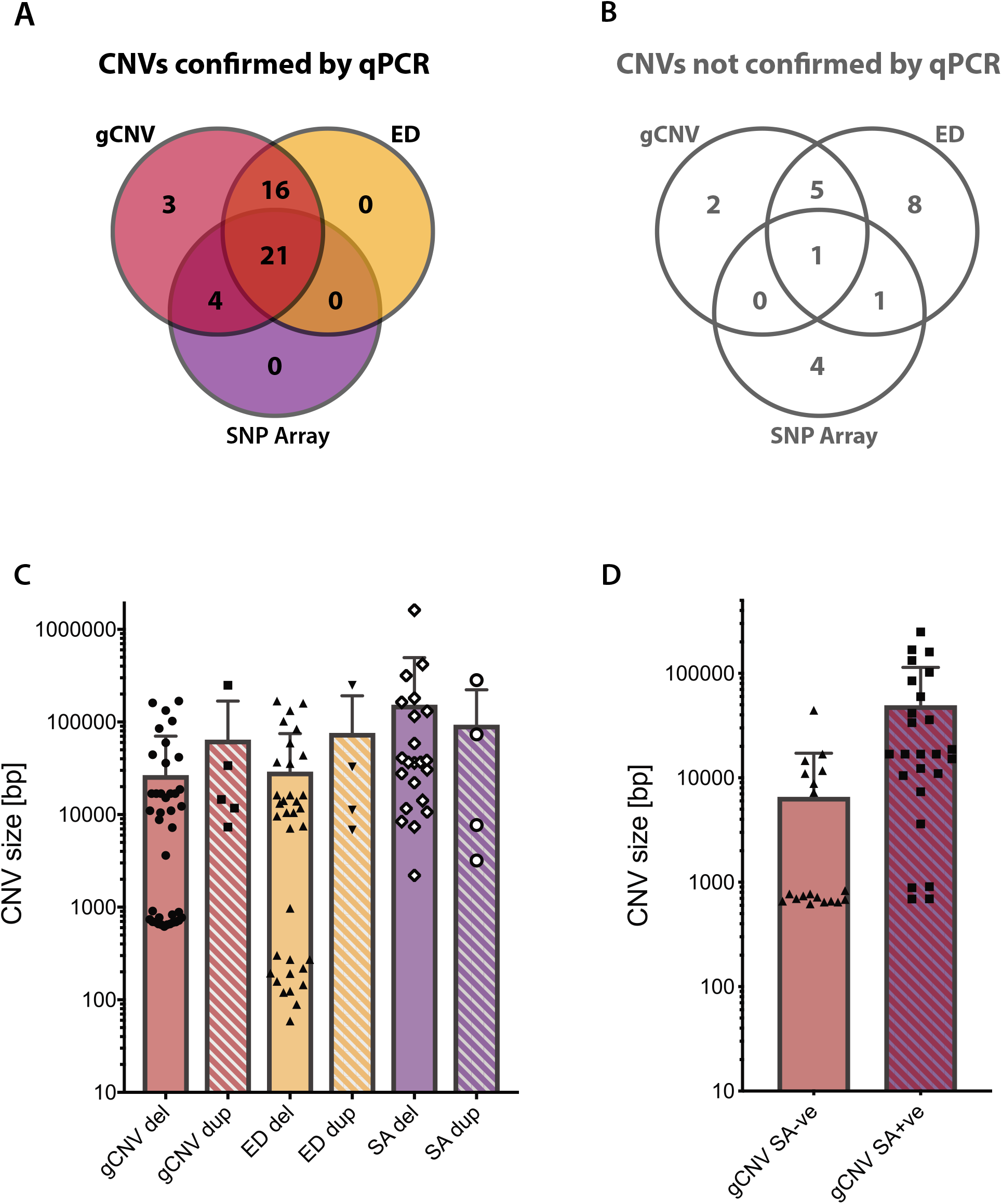
Comparison of Different CNV Detection Methods: A) Breakdown of true positive CNVs predicted by different methods. Note that all true positive CNVs were predicted by gCNV. B) Breakdown of false positive CNVs predicted by different methods. Note that only one false positive CNV was predicted by all three CNV prediction methods. C) The distribution of sizes of CNVs predicted by different methods. Note that in general, duplications were predicted less often, and were on average larger in size than predicted deletions. D) Comparison of the gCNV predicted sizes of validated CNVs that were also predicted by the SNP array (gCNV SA+ve) versus the gCNV predicted size of validated CNVs that were not predicted by the SA (gCNV SAve).

**Table 1.**
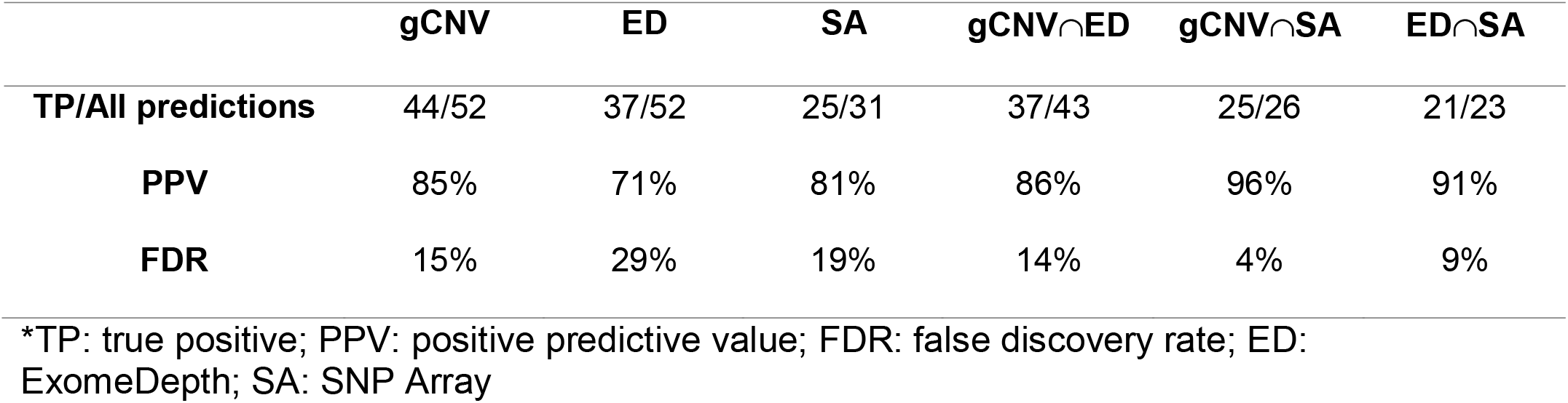
Positive predictive values for CNV detection

Depending on the technique used, the predicted sizes of the validated CNVs ranged from 59bp to 249kb (ExomeDepth), 619bp to 249kb (gCNV) and 2.2kb to1.6Mb (SNP-array) (Figure 2C). Since NGS-based methods only investigate regions covered by the capture kit (i.e. IRD gene exons), and the SNP-Array covers the genome in a more uniform fashion, the sizes of large CNVs predicted are probably more accurately represented by the SNP-Array. In this study, we did not attempt to map all the CNV breakpoints, however we validated the CNVs with multiple qPCR primers inside and outside of the predicted deletions (Supplementary Figure S1).

Further, we wanted to investigate the reasons for the SNP-array failing to detect 19 true CNVs that were predicted by gCNV. The most plausible reason is that the CNV is not covered by the sufficient number of SNPs on the array because it is too small or it is in a region that is poorly represented on the array. First, we compared the sizes of the CNVs predicted by gCNV that were validated by the SNP-Array (SA+ve) and that were not validated by the SNP-Array (SNP-ve) (Figure 2D). As expected CNVs that were not predicted by the SNP-Array were on average smaller (median = 772bp, mean = 6.5kb) than the CNVs that were also validated by the array (median = 17kb, mean = 49kb). None of the twelve CNVs that were below 1kb had sufficient number of SNPs (at least three) on the Omni2.5 Array to be detected by this method. Of the remaining seven larger CNVs (~7 - 44kb) that were not detected by the SNP-Array, most had sufficient number of SNPs in the interval (Supplementary Table S3), therefore, failure to detect them by the SNP array was due to other undetermined experimental or analytical reasons.

### Assessment of gCNV prediction scores

In the original analysis of the gCNV output we did not apply any quality filters, as the primary constraint on the data was whether a CNV fulfilled our “potentially solving” criteria. As such, we noticed that some low-quality predictions validated by qPCR. However, an agnostic search for true positive CNVs requires quality filters to remove false predictions, even at the expense of throwing out true positives. Therefore, we removed CNV predictions that were present in greater than 15% samples and CNVs from the opsin gene locus on chromosome X, which due to poor NGS mapping quality generated a high rate of likely false positive CNVs. Analysis of the 500 sample cohort with this filtering yielded 423 CNVs detected in 152 patients, of which 44 patients were solved with a CNV (validated by qPCR), 75 were solved with an SNV or small indel and 37 patients remained unsolved. Next, using the qPCR validated CNV predictions, we compared the different quality score metrics generated by gCNV (Figure 3) in order to choose one for a quality score cut-off. We settled on using the score of QA > 30^51^. This cut-off included three of nine falsely predicted CNVs and missed three of 47 true positive predictions (PPV=0.93) (Figure 3).

**Figure 3:**
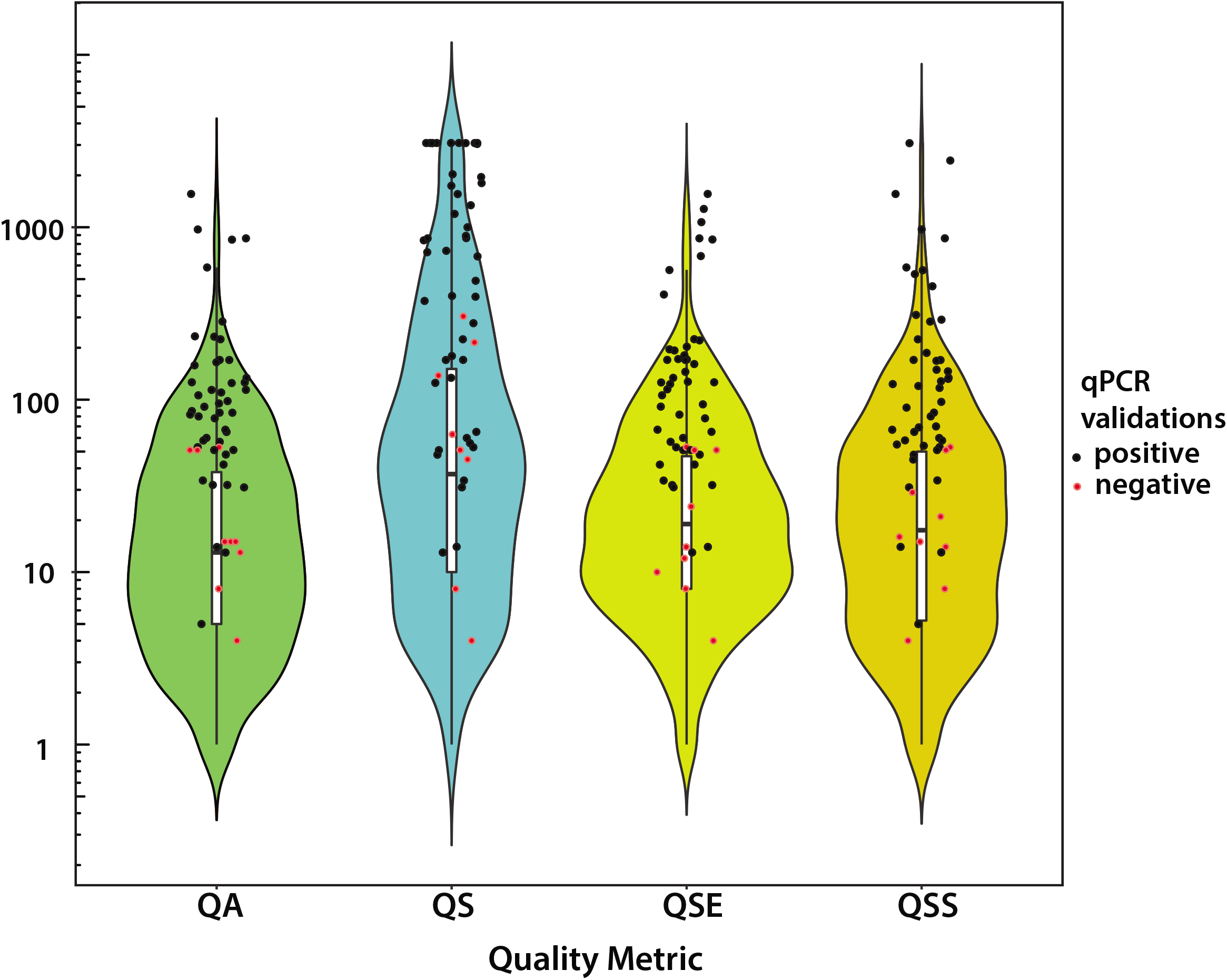
Distribution of gCNV Quality Metrics for Rare, Non-Opsin CNVs. The gCNV algorithm provides four quality metrics for every predicted CNV: QA, QS, QSE, and QSS. QA is the complementary Phred-scaled probability that **all points** (i.e. targets or bins) in the segment agree with the segment copy-number call. QS is the complementary Phred-scaled probability that at least one point (i.e. target or bin) in the segment agrees with the segment copy-number call. QSE is the complementary Phred-scaled probability that the segment end position is a genuine copy-number changepoint. QSS is the complementary Phred-scaled probability that the segment start position is a genuine copy-number changepoint. Violin plots show the relative probability density of the distribution of each quality metric, while internal box plots show the 25^th^, median, and 75^th^ percentile of distribution. The red and black data points represent CNVs within the distribution that were experimentally determined to be either true positives (black) or false positives (red) by qPCR validation.

### Assessment of frequency of patients with single likely pathogenic alleles in a recessive IRD gene

To assess how likely it is that the remaining genetically unsolved patients carry elusive pathogenic variants in already known IRD genes, we first evaluated how many unsolved patients have known pathogenic variants or new loss of function variants in recessive IRD disease genes, including CNVs. Of the 170 unsolved subjects, 38 carried stop, frameshift or essential splice-site variants and four subjects carried known pathogenic missense alleles. In addition, ten patients carried likely pathogenic CNVs (Supplementary Table S4). Altogether, we estimated that at least 30.6% of the unsolved patients (10.4% of the overall cohort) carried a single likely pathogenic allele in a recessive IRD gene.

## DISCUSSION

The results reported here indicate that CNVs contribute significantly to the genetic causality of IRDs, and that NGS based CNV detection methods outperform SNP array based CNV detection methods. Of the 500 patients whose genetic cause of disease was investigated by panel based NGS testing (GEDi)^66^, likely disease-causing CNVs were identified in 44 or 8.8% of cases. In 279 cases (55.8%) the disease could be explained by the likely pathogenic SNVs and small indels that are generated by the standard NGS analysis pipeline^51^. The only other structural variant that was investigated was a known *Alu* insertion in exon 9 of MAK^54–56^, which was present in seven cases (1.4%), agreeing closely with the previously reported frequency of 1.2%^54^.

The majority of the CNVs were heterozygous or homozygous deletions, ranging from single exon deletions to whole gene deletions. Three large deletions in two genes *(MERTK* and *TRPM1)* were in regions prone to the non-alleleic homologous recombination (NAHR), which is the most likely mechanism of their occurrence^41^.

Analysis of all CNVs reported in IRD genes performed by Van Schil and colleagues indicated that gene size, followed by the number of LINE and LTR repeats is the biggest predictor for a gene to be prone to CNVs^67^. This analysis correlates well with our findings, where the largest IRD gene *EYS* and the 3^rd^ largest IRD gene *USH2A*, had nine and eight causal CNVs respectively. However, in the present study the most common gene to harbor CNVs was *PRPF31*, which is known to cause a dominant form of IRD through haploinsufficiency^61^. A literature search by Van Schil and colleagues revealed that causal CNVs in *PRPF31* have been reported in 14 families to date, compared to 10 probands in the present study, indicating that CNVs in this gene were likely underestimated in the past. The reasons for such high frequency of CNVs in *PRPF31* is also unclear, since this gene is neither large (~16.3kb genomic length) nor does it have a high density of LINE and LTR repeats (ranked 191 of 245 genes by Van Schil and colleagues)^67^.

Searching for genetic causes of Mendelian diseases is commonly performed via a serial approach: first by looking for causal SNVs and small indels, and second moving on to more complex analyses such as CNV predictions. We propose that such causality searches are better undertaken in tandem because certain variants may falsely appear as homozygous but in fact they are *in trans* with a large deletion. This distinction is particularly important when a hypomorphic variant is involved, as in the case of p.Asn1868Ile in *ABCA4* discovered *in trans* with a large deletion in subject OGI655_001331. The p.Asn1868Ile variant in a homozygous state is not considered as pathogenic, however when paired *in trans* with a severe pathogenic variant it has been shown to be causal^62^. Therefore, only when considered together, the heterozygous deletion of nearly the entire gene with a heterozygous p.Asn1868Ile variant led to a conclusive genetic diagnosis.

Of 44 patients with CNVs, only five carried likely causal duplications, which may be a bias based on the fact that duplications are more difficult to detect, interpret and validate. All but one of the duplications is thought to result in a loss of function allele. Duplication of *OFD1* exons 6-15, is predicted to duplicate an internal 414 amino-acids of the protein and lead to a partially functioning OFD1 protein, resulting in a decreased spectrum of disease in subject OGI2829_004414, who apart from retinal disease has possible cognitive dysfunction and a history of renal failure, which resulted in transplantation. Variants in *OFD1* may lead to a spectrum of phenotypes from an x-linked dominant oral-facial-digital type 1 syndrome, with ciliopathy phenotype in females and embryonic lethal in males, x-linked recessive Joubert syndrome to non-syndromic IRD, depending on where the variant is located^39,65,68^, however to our knowledge no causal duplications in *OFD1* have been reported to date.

NGS based CNV prediction with the gCNV algorithm showed considerable advantage over more traditional SNP-array based prediction, as it had an increased diagnostic rate (44 vs 25 validated CNVs) and a higher positive predictive value (85% vs 81%). The major reason for this is that on average, NGS based algorithms could detect smaller CNVs as they were not restricted by the availability of SNPs in that region. In our study, all of the validated CNVs detected by the SNP array were also detected by the gCNV algorithm, therefore this method is an adequate replacement of the SNP-array based CNV predictions. Another accurate method of CNV detection is microarray-based comparative genomic hybridization (Array-CGH), which can be designed to cover intronic and exonic regions, as in the case of the IRD custom array (arrEYE)^47^. However, this method requires an additional wet-lab assay to be applied to the samples that had already undergone NGS, which may be unnecessary if the CNVs can be detected by a robust NGS-based algorithm. In this study we chose to use qPCR on genomic DNA as a validation of the CNVs, as this is a cost-effective and easily accessible method widely applied in many labs, however other assays such as Multiplex Ligation-dependent Probe Amplification (MLPA)^69^ or droplet PCR^70^ can also be used. Accurate characterization of CNVs can also be achieved by targeted locus amplification^67^.

In a genetic diagnostic setting, filtering based on CNV prevalence in the cohort (high prevalence could indicate likely capture artifact, or common CNV), CNV frequency per sample (high frequency could indicate low gDNA quality) and gCNV quality scores will aid in assessing the likelihood of a given CNV being true positive. In this study, we used experimental results to establish hard thresholds on gCNV predictions (discarding CNVs that appeared in >15% of the cohort, discarding CNVs with a QA < 30), to reduce predicted CNVs to a subset with a higher probability of being true genetic variants. In future studies, more sophisticated methods can be used, such as creating a scoring method which considers multiple factors, similar to the guidelines recommended for sequence variant curation from ACMG^71^. Taking into account the population-level frequency of CNVs using publicly available datasets will also be crucial in establishing the pathogenicity of the CNVs^67,72^.

In conclusion, our rate of discovery for likely solving variants in IRD patients has increased from 55.8% to 65.8% by including information from CNVs and MAK-Alu insertions. This represents a significant improvement in solving genetic cases, consistent with or higher than in previous studies^47,48,73–75^. Additional analysis found 8.4% of the cohort had a single potentially pathogenic SNV in a known autosomal recessive IRD gene, and 2% of the cohort had a single potentially pathogenic CNV in a known autosomal recessive IRD gene. Although this does not constitute enough evidence to identify that gene as disease causing, it indicates that there are high chances that a more elusive pathogenic variant resides on the second allele, as it has been demonstrated before for the *RPGRIP1* gene^76^. Searching for these elusive variants is critical for improving the discovery rate of disease causation. For example, this study did not investigate other SVs such as inversions or translocations, which can be difficult to detect outside of whole genome sequencing. We also looked at only one mobile element insertion, the well-reported MAK-Alu insertion. However, there is evidence that other mobile elements may play a role in IRDs^77^ and these could be searched for using bioinformatic software such as Mobster, which detects non-reference mobile element insertions in NGS data^78^. There have also been many examples of deep intronic variants leading to splicing aberration and retinal disease splicing variants can contribute to IRDs^39,42,44,76^, which are not readily available from NGS panel studies. Finally, variants in promoters, 3’/5’ UTRs, and other regulatory regions could contribute to disease etiologies^79^. The development of assays that can confirm pathogenic contributions from such variants, and the inclusion of such assays into variant analysis pipelines, will be important for better understanding the genetic contributions not just to IRDs, but to Mendelian disorders in general.

## Acknowledgments

This work was supported by grants from the National Eye Institute [R01EY012910 (EAP), R01EY026904 (KMB/EAP) and P30EY014104 (MEEI core support)], and the Foundation Fighting Blindness (EGI-GE-1218-0753-UCSD, KMB/EAP). The authors would like to thank the patients and their family members for their participation in this study and the Ocular Genomics Institute Genomics Core members for their experimental assistance. The authors would like to thank the Exome Aggregation Consortium, the Genome Aggregation Database (GnomAD) and the groups that provided exome variant data for comparison. A full list of contributing groups can be found at http://exac.broadinstitute.org/about and http://gnomad.broadinstitute.org/about.

## REFERENCES

1. Berger W, Kloeckener-Gruissem B, Neidhardt J. The molecular basis of human retinal and vitreoretinal diseases. Prog Retin Eye Res. 2010;29(5):335–375. doi:10.1016/j.preteyeres.2010.03.004

2. Hartong DT, Berson EL, Dryja TP. Retinitis pigmentosa. Lancet. 2006;368(9549): 1795–1809. doi:10.1016/S0140-6736(06)69740-7

3. Retinal Information Network. https://sph.uth.edu/retnet/home.htm. Published 2018. Accessed November 20, 2018.

4. Arno G, Agrawal SA, Eblimit A, et al. Mutations in REEP6 Cause Autosomal-Recessive Retinitis Pigmentosa. Am J Hum Genet. 2016;99:1–11. doi:10.1016/j.ajhg.2016.10.008

5. Agrawal SA, Burgoyne T, Eblimit A, et al. REEP6 deficiency leads to retinal degeneration through disruption of ER homeostasis and protein trafficking. Hum Mol Genet. 2017;26(14):2667–2677. doi:10.1093/hmg/ddx149

6. El Shamieh S, Neuillé M, Terray A, et al. Whole-exome sequencing identifies KIZ as a ciliary gene associated with autosomal-recessive rod-cone dystrophy. Am J Hum Genet. 2014;94(4):625–633. doi:10.1016/j.ajhg.2014.03.005

7. El-Asrag ME, Sergouniotis PI, McKibbin M, et al. Biallelic mutations in the autophagy regulator DRAM2 cause retinal dystrophy with early macular involvement. Am J Hum Genet. 2015;96(6):948–954. doi:10.1016/j.ajhg.2015.04.006

8. Goyal S, Jäger M, Robinson PN, Vanita V. Confirmation of TTC8 as a disease gene for nonsyndromic autosomal recessive retinitis pigmentosa (RP51). Clin Genet. July 2015. doi:10.1111/cge.12644

9. Ma X, Guan L, Wu W, et al. Whole-exome sequencing identifies OR2W3 mutation as a cause of autosomal dominant retinitis pigmentosa. Sci Rep. 2015;5:9236. doi:10.1038/srep09236

10. Audo I, Bujakowska K, Orhan E, et al. The familial dementia gene revisited: A missense mutation revealed by whole-exome sequencing identifies ITM2B as a candidate gene underlying a novel autosomal dominant retinal dystrophy in a large family. Hum Mol Genet. 2014;23(2). doi:10.1093/hmg/ddt439

11. Biswas P, Ramana V, Chavali M, et al. A missense mutation in ASRGL1 is involved in causing autosomal recessive retinal degeneration. Hum Mol Genet. 2016;25(12):2483–2497. doi:10.1093/hmg/ddw113

12. Avila-Fernandez A, Perez-Carro R, Corton M, et al. Whole-exome sequencing reveals ZNF408 as a new gene associated with autosomal recessive retinitis pigmentosa with vitreal alterations. Hum Mol Genet. 2015;24(14):4037–4048. doi:10.1093/hmg/ddv140

13. Coppieters F, Ascari G, Dannhausen K, et al. Isolated and Syndromic Retinal Dystrophy Caused by Biallelic Mutations in RCBTB1, a Gene Implicated in Ubiquitination. Am J Hum Genet. 2016;99(2):470–480. doi:10.1016/j.ajhg.2016.06.017

14. Audo I, Bujakowska K, Orhan E, et al. The familial dementia gene revisited: a missense mutation revealed by whole-exome sequencing identifies ITM2B as a candidate gene underlying a novel autosomal dominant retinal dystrophy in a large family. Hum Mol Genet. 2014;23(2):491–501. doi:10.1093/hmg/ddt439

15. Branham K, Matsui H, Biswas P, et al. Establishing the involvement of the novel gene *AGBL5* in retinitis pigmentosa by whole genome sequencing. Physiol Genomics. 2016;48(12):922–927. doi:10.1152/physiolgenomics.00101.2016

16. Wu JH, Liu JH, Ko YC, et al. Haploinsufficiency of RCBTB1 is associated with Coats disease and familial exudative vitreoretinopathy. Hum Mol Genet. 2016;25(8): 1637–1647. doi:10.1093/hmg/ddw041

17. Xu M, Eblimit A, Wang J, et al. ADIPOR1 Is Mutated in Syndromic Retinitis Pigmentosa. Hum Mutat. 2016;37(3):246–249. doi:10.1002/humu.22940

18. Zhang J, Wang C, Shen Y, et al. A mutation in ADIPOR1 causes nonsyndromic autosomal dominant retinitis pigmentosa. Hum Genet. 2016;135(12):1375–1387. doi:10.1007/s00439-016-1730-2

19. Saksens NTM, Krebs MP, Schoenmaker-Koller FE, et al. Mutations in CTNNA1 cause butterfly-shaped pigment dystrophy and perturbed retinal pigment epithelium integrity. Nat Genet. 2016;48(2): 144–151. doi:10.1038/ng.3474

20. Vincent A, Audo I, Tavares E, et al. Biallelic Mutations in GNB3 Cause a Unique Form of Autosomal-Recessive Congenital Stationary Night Blindness. Am J Hum Genet. 2016;98(5):1011–1019. doi:10.1016/j.ajhg.2016.03.021

21. Arno G, Holder GE, Chakarova C, et al. Recessive Retinopathy Consequent on Mutant G-Protein β Subunit 3 (GNB3). JAMA Ophthalmol. 2016;134(8):924–927. doi:10.1001/jamaophthalmol.2016.1543

22. Segarra NG, Ballhausen D, Crawford H, et al. NBAS mutations cause a multisystem disorder involving bone, connective tissue, liver, immune system, and retina. Am J Med Genet Part A. 2015;167(12):2902–2912. doi:10.1002/ajmg.a.37338

23. Bujakowska KM, Zhang Q, Siemiatkowska AM, et al. Mutations in IFT172 cause isolated retinal degeneration and Bardet-Biedl syndrome. Hum Mol Genet. 2015;24(1):230–242. doi:10.1093/hmg/ddu441

24. Consugar MB, Navarro-Gomez D, Place EM, et al. Panel-based genetic diagnostic testing for inherited eye diseases is highly accurate and reproducible, and more sensitive for variant detection, than exome sequencing. Genet Med. 2014; 17(4):253–261. doi:10.1038/gim.2014.172

25. Audo I, Bujakowska KM, Léveillard T, et al. Development and application of a next-generation-sequencing (NGS) approach to detect known and novel gene defects underlying retinal diseases. Orphanet J Rare Dis. 2012;7:8. doi:10.1186/1750-1172-7-8

26. Huang X-F, Huang F, Wu K-C, et al. Genotype-phenotype correlation and mutation spectrum in a large cohort of patients with inherited retinal dystrophy revealed by next-generation sequencing. Genet Med. 2015;17(4):271–278. doi:10.1038/gim.2014.138

27. Neveling K, Collin RWJ, Gilissen C, et al. Next-generation genetic testing for retinitis pigmentosa. Hum Mutat. 2012;33(6):963–972. doi:10.1002/humu.22045

28. Wang X, Wang H, Sun V, et al. Comprehensive molecular diagnosis of 179 Leber congenital amaurosis and juvenile retinitis pigmentosa patients by targeted next generation sequencing. J Med Genet. 2013;50(10):674–688. doi:10.1136/jmedgenet-2013-101558

29. Daiger SP, Bowne SJ, Sullivan LS, et al. Application of next-generation sequencing to identify genes and mutations causing autosomal dominant retinitis pigmentosa (adRP). Adv Exp Med Biol. 2014;801:123–129. doi:10.1007/978-1-4614-3209-8_16

30. Clark GR, Crowe P, Muszynska D, et al. Development of a diagnostic genetic test for simplex and autosomal recessive retinitis pigmentosa. Ophthalmology. 2010; 117(11):2169–2177. doi:10.1016/j.ophtha.2010.02.029

31. Bujakowska KM, Consugar MB, Place E, et al. Targeted exon sequencing in Usher syndrome type I. Invest Ophthalmol Vis Sci. 2014;55(12):8488–8496. doi:10.1167/iovs.14-15169

32. Glöckle N, Kohl S, Mohr J, et al. Panel-based next generation sequencing as a reliable and efficient technique to detect mutations in unselected patients with retinal dystrophies. Eur J Hum Genet. 2014;22(1):99–104. doi:10.1038/ejhg.2013.72

33. Weisschuh N, Mayer AK, Strom TM, et al. Mutation detection in patients with retinal dystrophies using targeted next generation sequencing. PLoS One. 2016;11(1):1–15. doi:10.1371/journal.pone.0145951

34. Zhao L, Wang F, Wang H, et al. Next-generation sequencing-based molecular diagnosis of 82 retinitis pigmentosa probands from Northern Ireland. Hum Genet. 2015; 134(2):217–230. doi:10.1007/s00439-014-1512-7

35. Boulanger-Scemama E, El Shamieh S, Démontant V, et al. Next-generation sequencing applied to a large French cone and cone-rod dystrophy cohort: Mutation spectrum and new genotype-phenotype correlation. Orphanet J Rare Dis. 2015; 10(1). doi:10.1186/s13023-015-0300-3

36. Perez-Carro R, Corton M, Sánchez-Navarro I, et al. Panel-based NGS Reveals Novel Pathogenic Mutations in Autosomal Recessive Retinitis Pigmentosa. Sci Rep. 2016;6(December 2015):1–10. doi:10.1038/srep19531

37. Bernardis I, Chiesi L, Tenedini E, et al. Unravelling the Complexity of Inherited Retinal Dystrophies Molecular Testing□: Added Value of Targeted Next-Generation Sequencing. Biomed Res Int. 2016;2016:6341870. doi:10.1155/2016/6341870

38. Costa KA, Salles MV, Whitebirch C, Chiang J, Sallum JMF. Gene panel sequencing in Brazilian patients with retinitis pigmentosa. Int J Retin Vitr. 2017;3(1):33. doi:10.1186/s40942-017-0087-6

39. Webb TR, Parfitt DA, Gardner JC, et al. Deep intronic mutation in OFD1, identified by targeted genomic next-generation sequencing, causes a severe form of X-linked retinitis pigmentosa (RP23). Hum Mol Genet. 2012;21(16):3647–3654. doi:10.1093/hmg/dds194

40. Braun TA, Mullins RF, Wagner AH, et al. Non-exomic and synonymous variants in ABCA4 are an important cause of Stargardt disease. Hum Mol Genet. 2013;22(25):5136–5145. doi:10.1093/hmg/ddt367

41. Bujakowska KM, Fernandez-Godino R, Place E, et al. Copy-number variation is an important contributor to the genetic causality of inherited retinal degenerations. Genet Med. 2017;19(6):643–651. doi:10.1038/gim.2016.158

42. Rio Frio T, McGee TL, Wade NM, et al. A single-base substitution within an intronic repetitive element causes dominant retinitis pigmentosa with reduced penetrance. Hum Mutat. 2009;30(9):1340–1347. doi:10.1002/humu.21071

43. Vaché C, Besnard T, le Berre P, et al. Usher syndrome type 2 caused by activation of an USH2A pseudoexon: implications for diagnosis and therapy. Hum Mutat. 2012;33(1):104–108. doi:10.1002/humu.21634

44. den Hollander AI, Koenekoop RK, Yzer S, et al. Mutations in the CEP290 (NPHP6) gene are a frequent cause of Leber congenital amaurosis. Am J Hum Genet. 2006;79(3):556–561. doi:10.1086/507318

45. Eisenberger T, Neuhaus C, Khan AO, et al. Increasing the yield in targeted next-generation sequencing by implicating CNV analysis, non-coding exons and the overall variant load: the example of retinal dystrophies. PLoS One. 2013;8(11):e78496. doi:10.1371/journal.pone.0078496

46. Pieras JI, Barragán I, Borrego S, et al. Copy-number variations in EYS: A significant event in the appearance of arRP. Investig Ophthalmol Vis Sci. 2011;52(8):5625–5631. doi:10.1167/iovs.11-7292

47. Van Cauwenbergh C, Van Schil K, Cannoodt R, et al. ArrEYE: A customized platform for high-resolution copy number analysis of coding and noncoding regions of known and candidate retinal dystrophy genes and retinal noncoding RNAs. Genet Med. 2017;19(4):457–466. doi:10.1038/gim.2016.119

48. Khateb S, Hanany M, Khalaileh A, et al. Identification of genomic deletions causing inherited retinal degenerations by coverage analysis of whole exome sequencing data. J Med Genet. 2016;53(9):600–607. doi:10.1136/jmedgenet-2016-103825

49. Neuhaus C, Eisenberger T, Decker C, et al. Next-generation sequencing reveals the mutational landscape of clinically diagnosed Usher syndrome: copy number variations, phenocopies, a predominant target for translational read-through, and PEX26 mutated in Heimler syndrome. Mol Genet Genomic Med. 2017;5(5):531–552. doi:10.1002/mgg3.312

50. Plagnol V, Curtis J, Epstein M, et al. A robust model for read count data in exome sequencing experiments and implications for copy number variant calling. Bioinformatics. 2012;28(21):2747–2754. doi:10.1093/bioinformatics/bts526

51. McKenna A, Hanna M, Banks E, et al. The Genome Analysis Toolkit: A MapReduce framework for analyzing next-generation DNA sequencing data. Genome Res. 2010;9(9):1297–1303. doi:10.1101/gr.107524.110.20

52. Babadi M. A Scalable, Bayesian Model For Copy Number Variation. https://www.broadinstitute.org/videos/scalable-bayesian-model-copy-number-variation-bayesian-pca.

53. Wang K, Li M, Hadley D, et al. PennCNV: An integrated hidden Markov model designed for high-resolution copy number variation detection in whole-genome SNP genotyping data. Genome Res. 2007;17(11):1665–1674. doi:10.1101/gr.6861907

54. Tucker B a, Scheetz TE, Mullins RF, et al. Exome sequencing and analysis of induced pluripotent stem cells identify the cilia-related gene male germ cell-associated kinase (MAK) as a cause of retinitis pigmentosa. Proc Natl Acad Sci U S A. 2011;108(34): E569–E576. doi:10.1073/pnas.1108918108

55. Özgül RK, Siemiatkowska AM, Yücel D, Myers CA, Collin RWJ, Zonneveld MN. Exome Sequencing and Cis-Regulatory Mapping Identify Mutations in MAK, Encoding a Regulator of Ciliary Length, as a Cause of Retinitis Pigmentosa Figure S1. Candidate Gene Prioritization at 34 Autosomal Recessive Retinitis Pigmentosa Loci. 89.

56. Bujakowska KM, White J, Place E, Consugar M, Comander J. Efficient in silico identification of a common insertion in the MAK gene which causes retinitis pigmentosa. PLoS One. 2015; 10(11). doi:10.1371/journal.pone.0142614

57. McLaren W, Gil L, Hunt SE, et al. The Ensembl Variant Effect Predictor. Genome Biol. 2016; 17(1). doi:10.1186/s13059-016-0974-4

58. Farkas M, Grant G, White J, Sousa M, Consugar M, Pierce E. Transcriptome analyses of the human retina identify unprecedented transcript diversity and 3.5 Mb of novel transcribed sequence via significant alternative splicing and novel genes. BMC Genomics. 2013;14(486).

59. Livak, Kenneth J, Schmittgen, Thomas D. Analysis of relative gene expression data using real-time quantitative PCR and the 2(-Delta Delta C(T)) Method. Methods. 2001. doi:10.1006/meth.2001.1262

60. Bujakowska KM, White J, Place E, Consugar M, Comander J. Efficient in silico identification of a common insertion in the MAK gene which causes retinitis pigmentosa. PLoS One. 2015; 10(11). doi:10.1371/journal.pone.0142614

61. Abu-Safieh L, Vithana EN, Mantel I, et al. A large deletion in the adRP gene PRPF31: evidence that haploinsufficiency is the cause of disease. Mol Vis. 2006;12:384–388. http://www.ncbi.nlm.nih.gov/entrez/query.fcgi?cmd=Retrieve&db=PubMed&dopt=Citation&list_uids=16636657.

62. Zernant J, Lee W, Collison FT, et al. Frequent hypomorphic alleles account for a significant fraction of ABCA4 disease and distinguish it from age-related macular degeneration. J Med Genet. 2017;54:404–412. doi:10.1136/jmedgenet-2017-104540

63. Oda T, Elkahloun AG, Pike BL, et al. Mutations in the human Jagged1 gene are responsible for Alagille syndrome. Nat Genet. 1997;16(July):235–242.

64. Kim BJ, Fulton AB. The genetics and ocular findings of Alagille syndrome. Semin Ophthalmol. 2007;22(4):205–210. doi:10.1080/08820530701745108

65. Ferrante MI, Giorgio G, Feather S a, et al. Identification of the gene for oral-facial-digital type I syndrome. Am J Hum Genet. 2001;68(3):569–576. http://www.pubmedcentral.nih.gov/articlerender.fcgi?artid=1274470&tool=pmcentrez&rendertype=abstract.

66. Consugar MB, Navarro-Gomez D, Place EM, et al. Panel-based genetic diagnostic testing for inherited eye diseases is highly accurate and reproducible, and more sensitive for variant detection, than exome sequencing. Genet Med. 2015; 17(4). doi:10.1038/gim.2014.172

67. Van Schil K, Naessens S, Van De Sompele S, et al. Mapping the genomic landscape of inherited retinal disease genes prioritizes genes prone to coding and noncoding copy-number variations. Genet Med. 2018;20(2):202–213. doi:10.1038/gim.2017.97

68. Coene KLM, Roepman R, Doherty D, et al. OFD1 is mutated in X-linked Joubert syndrome and interacts with LCA5-encoded lebercilin. Am J Hum Genet. 2009;85(4):465–481. doi:10.1016/j.ajhg.2009.09.002

69. Schouten JP, Mcelgunn CJ, Waaijer R, Zwijnenburg D, Diepvens F, Pals G. Relative quantification of 40 nucleic acid sequences by multiplex ligation-dependent probe amplification. Nucleic Acids Res. 2002;30(12):e57.

70. Hindson CM, Chevillet JR, Briggs HA, et al. Absolute quantification by droplet digital PCR versus analog real-time PCR. Nat Methods. 2013;10(10):1003–1005. doi:10.1038/nmeth.2633

71. Richards S, Aziz N, Bale S, et al. Standards and guidelines for the interpretation of sequence variants: a joint consensus recommendation of the American College of Medical Genetics and Genomics and the Association for Molecular Pathology. Genet Med. 2015;17(5):405–423. doi:10.1038/gim.2015.30

72. Collins RL, Brand H, Karczewski KJ, et al. An open resource of structural variation for medical and population genetics. bioR. 2019;March 14:1–15.

73. Huang X, Mao J, Huang Z, et al. Genome-Wide Detection of Copy Number Variations in Unsolved Inherited Retinal Disease. Investig Ophthalmol Vis Sci. 2017;58(1):424–429. doi:10.1167/iovs.16-20705

74. Ellingford JM, Horn B, Campbell C, et al. Assessment of the incorporation of CNV surveillance into gene panel next-generation sequencing testing for inherited retinal diseases. J Med Genet. 2018;55:114–121. doi:10.1136/jmedgenet-2017-104791

75. Jespersgaard C, Fang M, Bertelsen M, et al. Molecular genetic analysis using targeted NGS analysis of 677 individuals with retinal dystrophy. Sci Rep. 2019;9(1219): 1–7. doi:10.1038/s41598-018-38007-2

76. Jamshidi F, Place EM, Mehrotra S, et al. Contribution of non-coding mutations to RPGRIP1-mediated inherited retinal degeneration. Genet Med. 2018;ePub(August 03): 1–11. doi:doi.org/10.1101/211292

77. Tavares E, Tang CY, Li S, et al. Retrotransposon insertion as a novel mutational event in Bardet - Biedl syndrome. Mol Genet Genomic Med. 2019;7(2):e521. doi:10.1002/mgg3.521

78. Thung DT jwa., de Ligt J, Vissers LEM, et al. Mobster: accurate detection of mobile element insertions in next generation sequencing data. Genome Biol. 2014. doi:10.1186/s13059-014-0488-x

79. Coppieters F, Todeschini AL, Fujimaki T, et al. Hidden Genetic Variation in LCA9-Associated Congenital Blindness Explained by 5’UTR Mutations and Copy Number Variations Of NMNAT1. Hum Mutat. 2015;12:1188–1196. doi:10.1002/humu.22899

