## Supplementary Files for "Copy-number variation contributes 9% of pathogenicity in the inherited retinal degenerations"

**Methods:**

**RPGR ORF15 sequencing**

Exon 15 of *RPGR* transcript NM_001034853 (called *RPGR* ORF15) was PCR amplified with forward 5’-AGCCAGACAGTTACATGGAAGGTGCAA-3’ and reverse 5’-TGTCTTTGGCTCCTTAACACAGCTGCA-3’ primers ^1^ using TaKaRa LA Taq DNA Polymerase Hot-Start Version (Takara Bio USA, Inc., Mountain View, CA 94043 USA). The PCR product was purified (ExoSAP-IT PCR Product Cleanup Reagent, ThermoFisher Scientific, USA) and sequenced with a Sanger sequencing kit (BigDye™ Terminator v3.1 Cycle Sequencing Kit, ThermoFisher) with addition of 1x Solution S (SolisBiodyne, Mango Biotechnology, LLC, Mountain View, CA 94040, USA) in six separate reactions with the following primers:

R4: 5’-CTCTCCTTCCTCCTTTTCAC-3’

R7: 5’-CCTTCCTCCTCTTCCCCCTCA-3’

R8: 5’-TCCTTCCTCCTCTTCCCCCTCCCA-3’

R9: 5’-CCCTGTGTGTTAGTAACTGAC-3’

F10: 5’-GGAAATGGGAAAGAGCAGAGGTC-3’

R5: 5’-ACTGGCCATAATCGGGTCACAT-3’

The annealing and extension temperatures in the sequencing reaction were adjusted as follows: for primers R4, R5, R9, F10 (annealing 56°C, extension 60°C), primer R7 (annealing 58°C, extension 63°C), primer R8 (annealing 63°C, extension 68°C)^2^. The sequencing reactions were analyzed with a 96-capillary Sanger sequencer (ABI 3730xl, Applied Biosystems).

**ACMG Variant Classification:**

Potentially solving variants were classified according to ACMG guidelines^3^ (Supplementary Table S2). In 99 of the subjects, using the ACMG guidelines resulted in classifying one or more variant as a Variant of Uncertain Significance (VUS). In 52 of these cases, we still strongly suspect that the variant may be contributing to IRD, as it pairs with either a CNV, a Likely Pathogenic (LP) variant, or a Pathogenic (P) variant in the same gene. In the other 47 subjects, we found either two VUSes in a known ar IRD gene, or one VUS in a known ad or x-linked IRD gene. Because many of these variants do not have segregation data available, and functional studies are outside the scope of this report, more evidence could not be provided to confidently alter the variant status.

**Figure S1A**

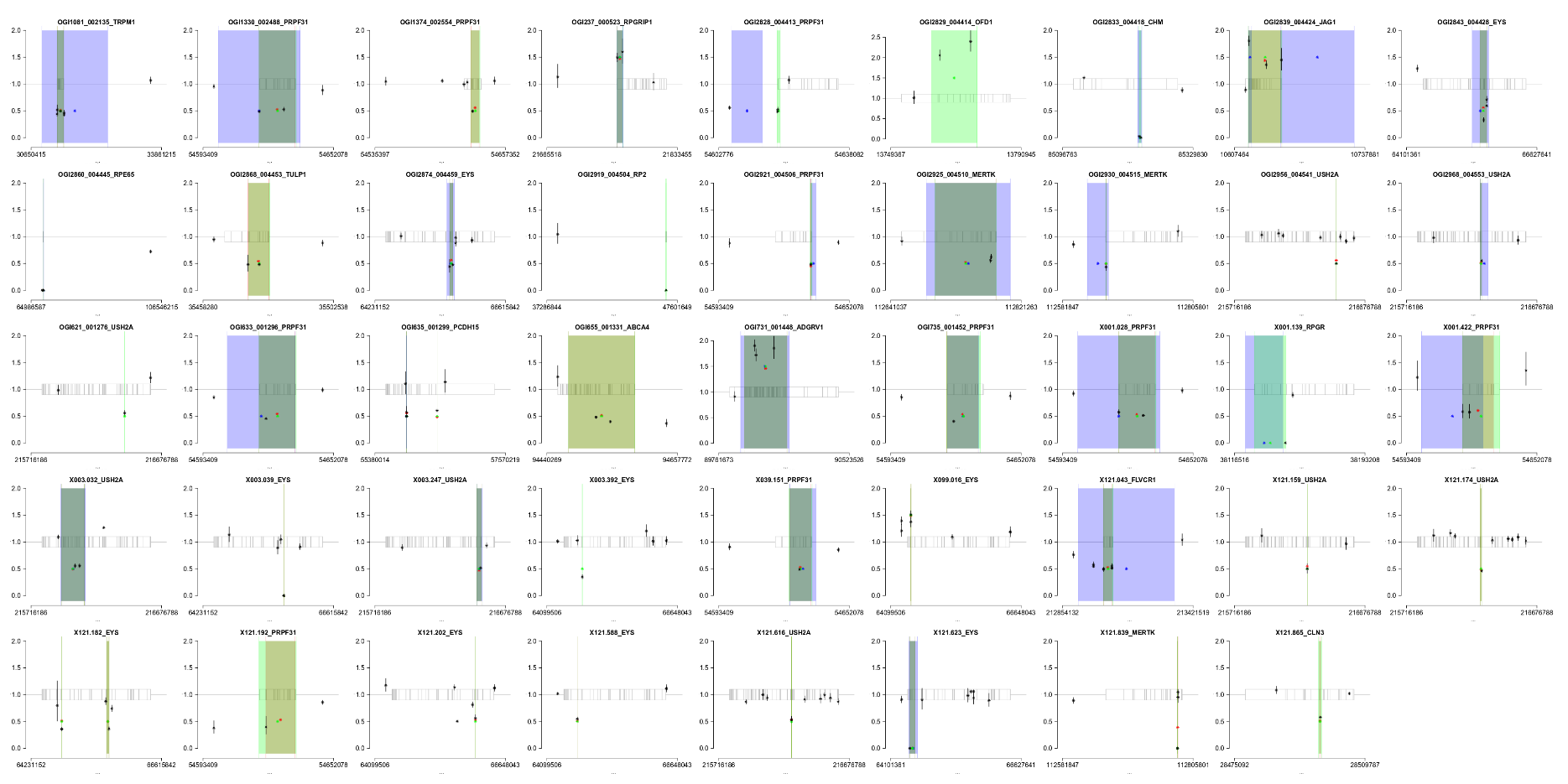

**Figure S1B**

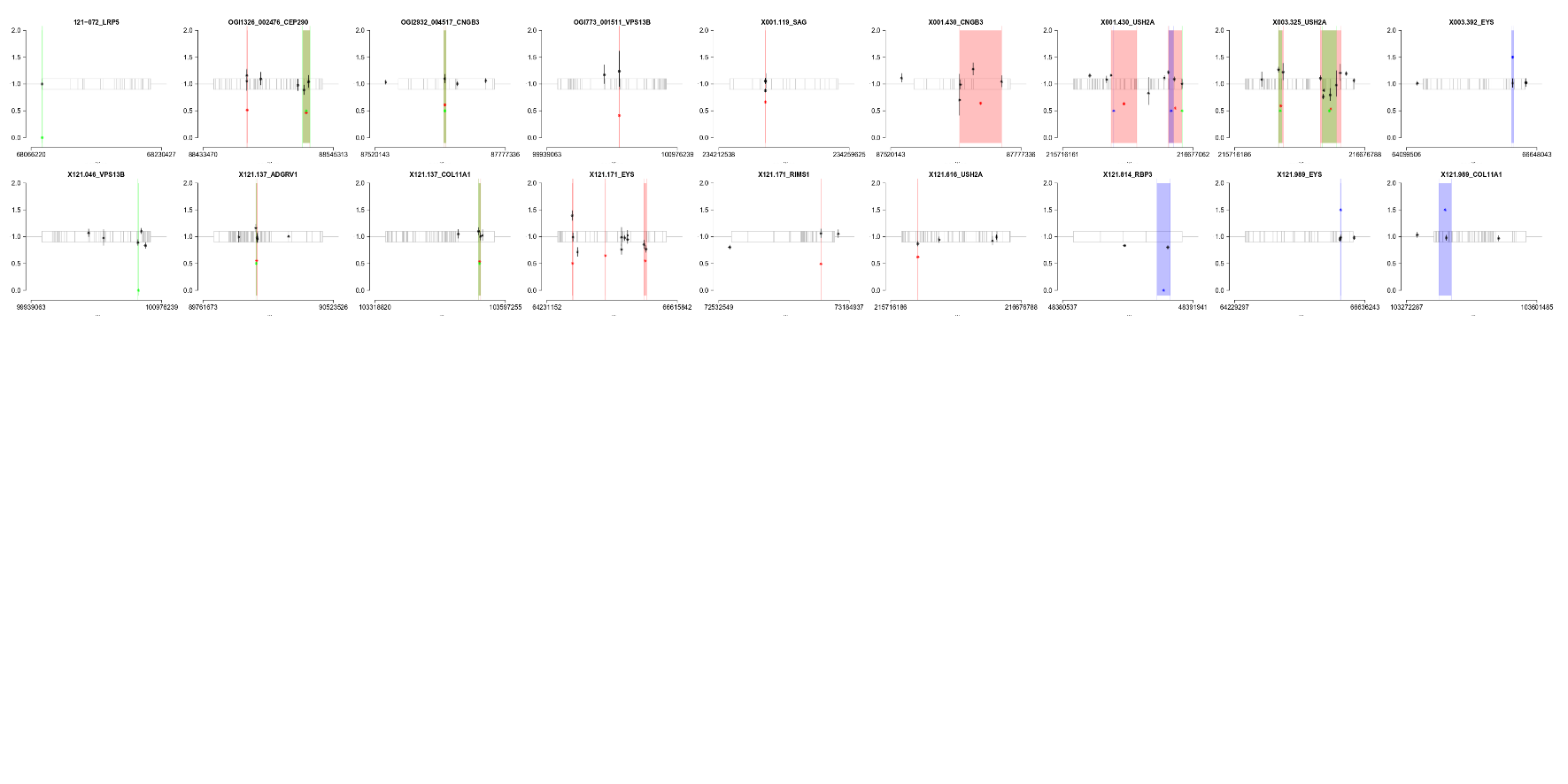

**Figure S1: qPCR validation of predicted CNVs.** Each graph represents a unique sample/gene combination that had one or more predicted CNVs by one or more methods. Red regions are CNVs predicted by ExomeDepth. Blue regions are CNVs predicted by SNP Array. Green regions are CNVs predicted by gCNV. The grey box with horizontal bars represents the gene and exons. Red/blue/green points indicate the fold change predictions of the various software. Black points indicate the experimentally determined fold change of the sample’s DNA at those genomic coordinates. Note that when there is a long distance between a predicted CNV and a flanking black point, this was because appropriate qPCR primers could not be found closer to the gene. A) Collection of all CNVs that were validated as true positives by qPCR. The CNVs in sample OGI2919_004504, gene RP2 and sample OGI655_001331, gene ABCA4 were not included in the analysis for Figure 2A, as the samples were never run on a SNP array. B) Collection of all CNVs that were validated as false positives by qPCR. The two CNVs in sample OGI1326_002476, gene CEP290 were not included in the analysis for Figure 2B, as the sample was never run on a SNP array. Note that for sample 121-072, gene LRP5, the CNV was validated with traditional PCR rather than qPCR.

**Supplementary References:**

1.	Li J, Tang J, Feng Y, et al. Improved Diagnosis of Inherited Retinal Dystrophies by High-Fidelity PCR of ORF15 followed by Next-Generation Sequencing. *J Mol Diagnostics*. 2016;18(6):817-824. doi:10.1016/j.jmoldx.2016.06.007

2.	Neidhardt J, Glaus E, Lorenz B, et al. Identification of novel mutations in X-linked retinitis pigmentosa families and implications for diagnostic testing. *Mol Vis*. 2008;14(January):1081-1093.

3.	Richards S, Aziz N, Bale S, et al. Standards and guidelines for the interpretation of sequence variants: a joint consensus recommendation of the American College of Medical Genetics and Genomics and the Association for Molecular Pathology. *Genet Med*. 2015;17(5):405-423. doi:10.1038/gim.2015.30
